# Metabolic switching and cell wall remodelling of *Mycobacterium tuberculosis* during bone tuberculosis

**DOI:** 10.1101/2022.01.13.476285

**Authors:** Khushpreet Kaur, Sumedha Sharma, Sudhanshu Abhishek Sinha, Prabhdeep Kaur, Uttam Chand Saini, Mandeep Singh Dhillon, Petros C. Karakousis, Indu Verma

**Author notes:** **Correspondence:** Prof. Indu Verma. These authors contributed equally to this work (Sumedha Sharma, Sudhanshu Abhishek Sinha).

## Abstract

Bone tuberculosis is widely characterized by irreversible bone destruction caused by *Mycobacterium tuberculosis*. Mycobacterium has the ability to adapt to various environmental stresses by altering its transcriptome in order to establish infection in the host. Thus, it is of critical importance to understand the transcriptional profile of *M. tuberculosis* during infection in the bone environment compared to axenic cultures of exponentially growing M.tb. In the current study, we characterized the in vivo transcriptome of *M. tuberculosis* within abscesses or necrotic specimens obtained from patients with bone TB using whole genome microarrays in order to gain insight into the *M. tuberculosis* adaptive response within this host microenvironment. A total of 914 mycobacterial genes were found to be significantly over-expressed and 1688 were repressed (fold change>2; *p*-value ≤0.05) in human bone TB specimens. Overall, the mycobacteria displayed a hypo-metabolic state with significant (*p*≤0.05) downregulation of major pathways involved in translational machinery, cellular and protein metabolism and response to hypoxia. However, significant enrichment (*p* ≤0.05) of amino-sugar metabolic processes, membrane glycolipid biosynthesis, amino acid biosynthesis (serine, glycine, arginine and cysteine) and accumulation of mycolyl-arabinogalactan-peptidoglycan complex suggests possible mycobacterial survival strategies within the bone lesions by strengthening its cell wall and cellular integrity. Data were also screened for M.tb virulence proteins using Virulent Pred and VICM Pred tools, which revealed five genes (*Rv1046c, Rv1230c, DppD, PE_PGRS26* and *PE_PGRS43*) with a possible role in the pathogenesis of bone TB. Next, an osteoblast cell line model for bone TB was developed allowing for significant intracellular multiplication of M.tb. Interestingly, three virulence genes (*Rv1046c, DppD* and *PE_PGRS26*) identified from human bone TB microarray data were also found to be overexpressed by intracellular *M. tuberculosis* in osteoblast cell lines. Overall, these data demonstrate that *M. tuberculosis* alters its transcriptome as an adaptive strategy to survive in the host and establish infection in bone. Additionally, the *in vitro* osteoblast model we describe may facilitate our understanding of the pathogenesis of bone TB.

**Author Summary:** Musculoskeletal tuberculosis is the third most common manifestation of extra-pulmonary tuberculosis and massive bone destruction along with vertebral discs are one of the hallmarks of this disease. *Mycobacterium tuberculosis*, the causative agent, has the tremendous potential to adapt itself to different host environments due to its ability to alter the expression of genes/proteins belonging to different pathways. This study shows that the mycobacterial infection in bone is driven by the increased expression of genes belonging to cell wall remodelling and DNA damage repair pathways important for its survival. Further data analysis showed that some of these genes are coding for proteins possessing virulence potential that may be essential for survival of *M. tuberculosis* under such hostile environment of bone. We also developed an in vitro model of bone tuberculosis using an osteoblast cell line and validated the expression of these virulence factors. Identification of such virulence factors in the bone environment by *M. tuberculosis* may aid to identify new therapeutic targets for bone TB. Further, development of cell line model for bone TB is important to understand some unknown facets of this disease.

## Introduction

Bone tuberculosis (TB) is one of the ancient extra-pulmonary manifestations of TB, as evidenced by the traces of DNA and cell wall mycolates detected in femurs of Egyptian mummies^1^. In India, it accounts for 10-25% of extra-pulmonary TB (EPTB) cases^2^ and the prevalence of disease has been increasing in other endemic countries. The spine (tuberculous spondylitis/Pott’s spine) is the most commonly affected skeletal site, accounting for >50% of bone TB cases. The disease is widely characterized by bone resorption, destruction of the vertebral bodies, discs, and formation of abscesses, eventually leading to vertebral collapse and kyphotic deformities^3^. Necrotic caseation and cold abscesses are the characteristic findings of tuberculous spondylitis in the majority of bone lesions^4^. Early diagnosis of bone TB is challenging due to its pauci-bacillary nature, deep inaccessible lesions, non-specific symptoms and resemblance of the disease to various other bone diseases and infections. Diagnosis of the disease mainly relies on clinical and radiological findings. However, if undiagnosed, it eventually leads to paraplegia and neurological abnormalities associated with long-term morbidity^5^. A few studies have revealed how mycobacterial infection disrupts the bone homeostasis and favours enhanced osteoclastogenesis, ultimately causing bone destruction^6,7^.

*Mycobacterium tuberculosis*, an obligate aerobic bacterium, thrives best in oxygen-rich environments such as the lungs, but it can also survive and establish infection in contained osseous tissue^8^. This is possible due to *M. tuberculosis* adaptations to different stresses and host environments, primarily involving alterations in its gene expression profile. Various studies have revealed the differential expression patterns of mycobacteria isolated from different sites within the human host^9–11^. The transcriptional profile of mycobacteria isolated from the lung granulomatous region is different from that obtained from normal-appearing lungs in *M. tuberculosis*-infected individuals^12^. The altered gene expression of *M. tuberculosis* under various physiologically relevant stress conditions, e.g., hypoxia and nutrient starvation, is accompanied by a switch from active growth to a more stable dormant state characterized by reduced metabolism and cell wall thickening^13,14^. Thus, in the present study, we aimed to understand mycobacterial adaptations to the bone environment by investigating its transcriptome within abscesses or necrotic specimens harvested from patients with bone TB. As a further step in elucidating key mycobacterial virulence factors in bone TB, we developed an in vitro osteoblast model of *M. tuberculosis* infection and confirmed the gene expression patterns of selected virulent proteins.

## Results

### Transcriptional profile of *M. tuberculosis* in human bone lesions

The transcriptional profile of *M. tuberculosis* within bone specimens obtained from patients (n=5) was analyzed in comparison to exponentially growing *M. tuberculosis* H37Rv using whole genome microarrays. Differentially expressed genes (DEGs) were filtered with a fold change cut off of 2 (>2 or ≤-2) and a p-value ≤0.05 (Fig. 1a). Data for the same have been submitted as gene expression omnibus (GEO) dataset vide accession no. (GSE-165232). Analysis revealed 2602 DEGs (Table S2), of which 914 genes were upregulated and 1688 were downregulated. A representative heat map and hierarchical clustering analysis of the same is shown in Figure 1b. DEGs were classified into functional categories as per the tuberculist database (Table 1).

**Fig. 1:**
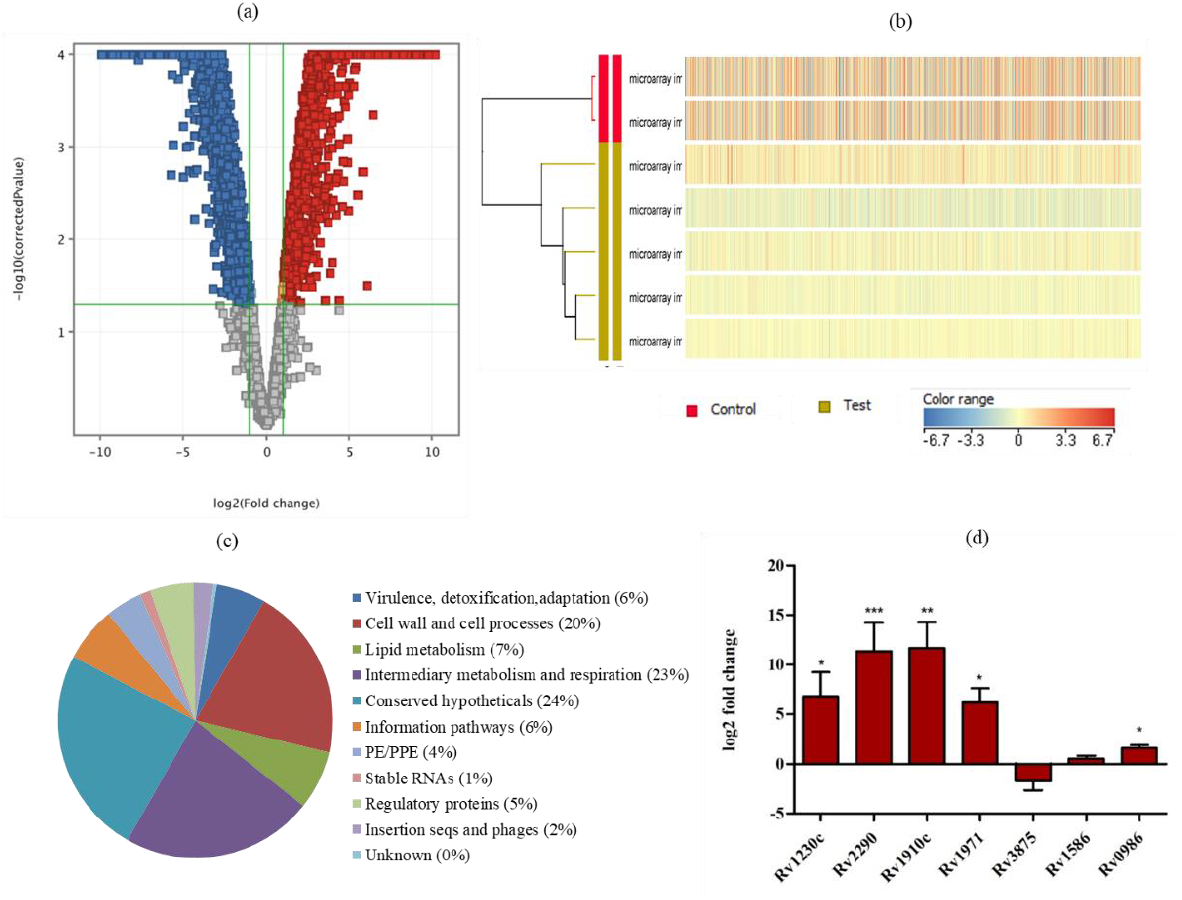
Transcriptional profile of *Mycobacterium tuberculosis* in abscess or necrotic tissue obtained from bone TB patients. (a) Volcano plot represents the distribution of all differentially expressed mycobacterial genes filtered on the basis of >2 or<-2 fold change and corrected p-value of 0.05. Red color represents upregulation, blue represents downregulation and grey with no change. (b) Heat map reflecting the hierarchical clustering of differentially expressed mycobacterial genes in bone TB specimen obtained from five different patients, red color represents upregulated genes, blue represents downregulated genes and yellow color with no change in gene expression. (c) Pie chart represents the functional categories of all differentially expressed mycobacterial genes as determined by Tuberculist (d) qRT-PCR validation of a subset of differentially expressed mycobacterial genes selected from microarray in human bone TB specimen (n=6).

**Table 1:** Functional categorization of differentially expressed genes of *Mycobacterium tuberculosis* in human bone TB specimens based on the tuberculist database.

| S. No. | Functional categories <sup>a</sup> | Upregulated | $p$-value<br>(Hypergeometric<br>Probability) | Downregulated | $p$-value<br>(Hypergeometric<br>Probability) | Total |
| --- | --- | --- | --- | --- | --- | --- |
| 1 | Virulence, detoxification, adaptation (239) <sup>b</sup> | 46 (5%) | **0.034** | 105 (6%) | **0.0348** | 151 |
| 2 | Cell wall and cell processes (772) | 183 (20%) | **0.021** | 347 (21%) | **0.0017** | 530 |
| 3 | Lipid metabolism (272) | 62 (7%) | 0.059 | 118 (7%) | **0.036** | 180 |
| 4 | Intermediary metabolism and respiration (936) | 213 (23%) | **0.032** | 374 (22%) | **0.022** | 587 |
| 5 | Conserved hypotheticals (1042) | 236 (26%) | **0.032** | 400 (24%) | **0.0037** | 636 |
| 6 | Information pathways (242) | 42 (5%) | **0.011** | 125 (7%) | **1.509 e-4** | 167 |
| 7 | PE/PPE (168) | 44 (5%) | **0.033** | 68 (4%) | 0.063 | 112 |
| 8 | Stable RNAs (80) | 2 (0%) | **3.986 e-7** | 31 (2%) | 0.0841 | 33 |
| 9 | Regulatory proteins (198) | 48 (5%) | 0.053 | 84 (5%) | 0.0542 | 132 |
| 10 | Insertion seqs and phages (147) | 35 (4%) | 0.070 | 29 (2%) | **1.264 e-8** | 64 |
| 11 | Unknown (15) | 3 (0%) | 0.245 | 7 (0%) | 0.184 | 10 |
| Total (4111) |  | 914 |  | 1688 |  | 2602 |

<sup>a</sup> Various functional Categories of *M. tuberculosis* H37Rv as listed on the tuberculist database
<sup>b</sup> Number represents the total number of genes in each tuberculist category of *M.tb* H37Rv.
<sup>c</sup> Bold values indicate statistical significance at $p < 0.05$

As shown in Figure 1c, the largest proportion of genes were classified as conserved hypothetical proteins (24%), followed by intermediary metabolism and respiration (23%), cell wall and cell processes (20%), lipid metabolism (7%) and virulence, detoxification, adaptations (6%) and information pathways (6%). A detailed analysis of upregulated and downregulated genes within each category is given in Table S3.

### Mycobacterial adaptation within the bone microenvironment

For further analysis of *M. tuberculosis* adaptation within the bone microenvironment, pathway enrichment analysis was performed using the Biocyc database and Fisher’s exact test with post hoc Benjamini-Hochberg correction (p≤0.05) for statistical analysis. This analysis identified significantly positively and negatively enriched pathways, transcriptional/translational regulators, and gene ontology (GO) terms among the upregulated and downregulated categories, respectively (Figs. 2a and 2b). Although there were no significant positively enriched pathways among upregulated DEGs after using post hoc correction, usage of only Fisher’s exact statistical analysis without post hoc correction led to identification of 77 positively enriched pathways with a p-value ≤ 0.05 (Table S4). Further, among downregulated DEGs, 38 negatively enriched pathways were observed (Table S5).

**Fig. 2:**
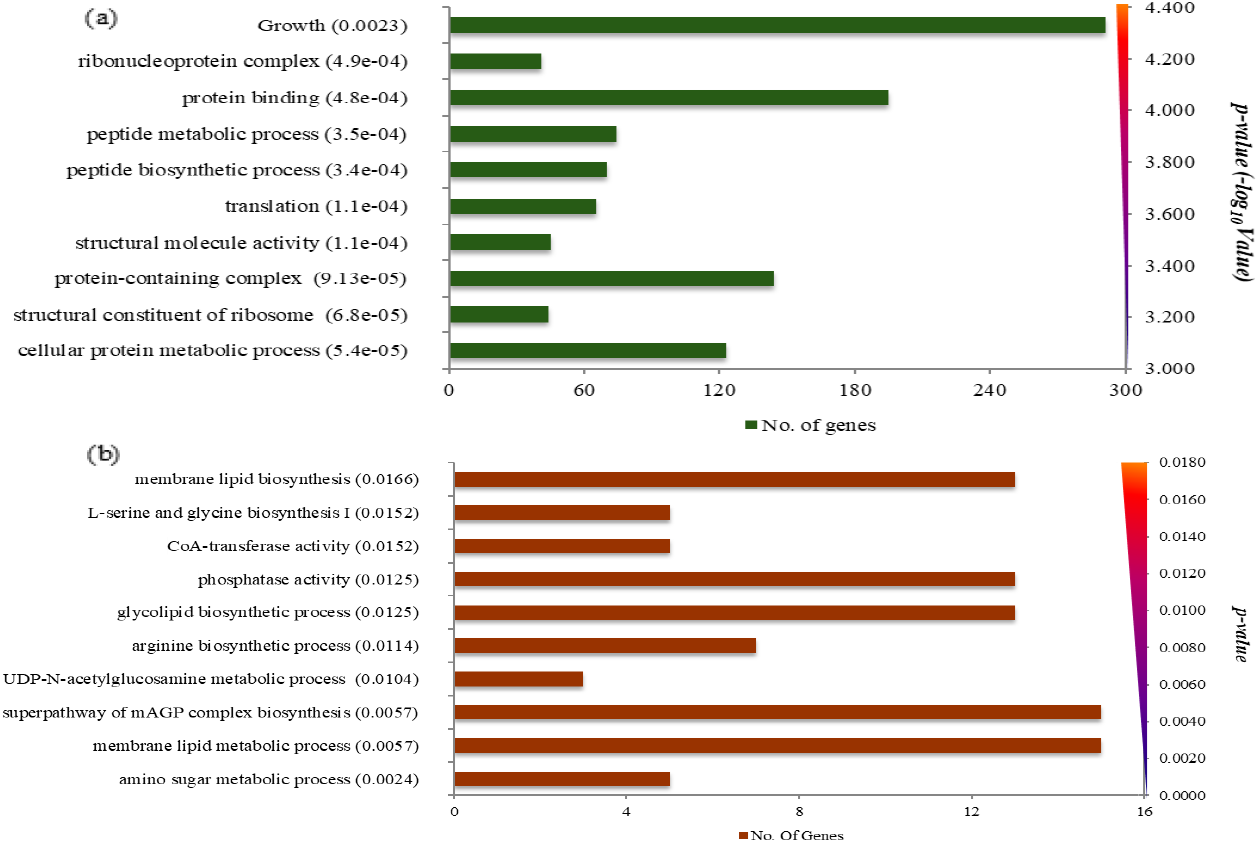
Significantly enriched pathways correspond to differentially upregulated and downregulated mycobacterial genes using Biocyc database. Bar graph for top 10 enriched pathways for differentially (a) downregulated pathways, *p-value* computed using fisher’s exact test and post-hoc Benjamini Hochberg’s correction. (b) upregulated pathways, *p*-value calculated using fisher’s exact test. X-axis shows the no. of genes and Y-axis represents p-value. *p-value* >0.05 is considered statistically significant.

### Upregulated pathways

Among the 77 positively enriched pathways, significantly enriched pathways belonged to amino-sugar metabolism (*p≤0.005*), membrane lipid metabolism and synthesis (*p≤0.005*), and biosynthesis of mycolyl-arabinogalactan-peptidoglycan (mAGP) complex (*p*≤0.005). Besides these, various other biosynthetic pathways were enriched, including synthesis of several amino acids (arginine, serine and glycine (*p*≤0.01)) and sulphate assimilation and cysteine biosynthesis (*p*≤0.01). Several regulatory pathways were also enriched, including protein phosphatases (p≤0.02) and cellular response to DNA damage stimulus (p≤0.02). Below we describe the details of the significantly enriched upregulated pathways using the Biocyc database^15^.

#### Cell wall structure and integrity

Peptidoglycans (PG), mycolic acid and arabinogalactan are major constituents of the mycobacterial cell wall and essential for bacterial survival under extreme conditions^16^. Genes involved in the synthesis of amino-sugar derivatives, like UDP-N-acetyl glucosamine (*glmU* (6.1), *glmS* (42.4), *mrsA* (2.69), *nagA* (5.4) and *Rv2267c* (5.3)), encoding a key metabolite and starting point for peptidoglycan synthesis^17^ were found to be overexpressed. Also, there was significant upregulation of genes involved in the formation of the mAGP complex, including (*glfT2 (90.5), embB (11.9), Ag85c (23.7), ubiA (6.9), Rv3807c (9.3), wecA (3.6), Rv2361c (13.2), Rv3468c (6.2), aftC (2.14), prsA (14.08), galE1(6.2)* and *galE3 (3.8)* etc). Additionally, the genes *Rv2174, Rv2181, pimB* and *Rv3631*, which encode enzymes involved in biosynthesis of membrane glycolipids, like lipomannan (LM)/ lipoarabinomannan (LAM), a major component in TB immune-pathogenesis, were also found to be upregulated. In addition, genes involved in the synthesis of phthiocerol-based lipids (cord factors) and glycolipids (*Rv2957, Rv2958c* and *Rv2962c*) were also upregulated. Two genes (*otsA, otsB2*), encoding enzymes involved in the synthesis of trehalose, a major structural constituent of cell wall glycolipids, were also found to be upregulated (Table S4).

#### Amino acid biosynthesis

Numerous genes of the amino acid biosynthetic pathways, including serine, glycine, cysteine and arginine, were induced in this study. Serine is an essential amino acid for mycobacterial growth^18^ and serves as a precursor for the synthesis of glycine, cysteine and phospholipids^19^. We observed significant induction of the genes *serB2* and *serA2*, which encode enzymes involved in serine biosynthesis. The gene *glyA2*, encoding serine hydroxyl methyl transferase, which converts serine to glycine, was also significantly upregulated. Furthermore, *M. tuberculosis* in bone lesions showed increased expression of the genes involved in sulphur accumulation and biosynthesis of cysteine, including *cys A1*, encoding a subunit of the sulphate transporter, and *cysN*, which encodes a subunit of ATP sulfurylase responsible for activating the imported sulphate to adenosine-5’-phosphosulfate (APS). The APS can further be used for sulfation of biomolecules or for the synthesis of reduced sulphur compounds, such as cysteine, which can be converted to methionine and mycothiol. Among the cysteine synthesis genes required to reduce APS to cysteine, the gene *cysK*, which encodes for O-acetylserine sulfhydrylase, was also upregulated. This gene codes for one of the enzymes required to condense sulphide with O-acetyl serine. As described above, several genes involved in the biosynthesis of serine, which forms the intermediate O-acety1 serine for cysteine synthesis, were also upregulated. In addition to serine, glycine and cysteine biosynthesis, multiple genes involved in the *de novo* arginine biosynthesis pathway (*argC, argD, argG, carB* and *carA*) were also upregulated in *M. tuberculosis* within human bone lesions (Table S3). Arginine biosynthesis is also an essential pathway and has been reported to be upregulated in response to oxidative stress in *M. tuberculosis*^20^ and also arginine deprivation cause mycobacterial cell death^21^.

#### Protein phosphatases (Regulators of cell processes and virulence)

Serine/threonine phosphatases are essential signalling enzymes in bacteria that catalyze the hydrolysis of some phospho-substrates which further control cell cycle events and intracellular survival of the bacteria^22^. Several genes encoding for enzymes possessing phosphatase activity (*PstP, Rv1364c, gpgP, pknG, Rv3807, Rv3376, Rv3813* and *Rv1225c)* were found to be upregulated in this study. Among them, *pstP*, a key regulator of phosphorylation and cell division^23^ and *pknG*, which senses the availability of amino acids under nutrient deprived conditions^24^ were also upregulated in bone TB lesions. In addition to these, genes encoding acid phosphatases (*Rv2577* and *Rv2135c*) were also found to be upregulated in the current study.

#### DNA damage response

Intracellular pathogens, such as *M. tuberculosis*, experience a variety of DNA-damaging assaults in vivo, including the oxidative burst and host responses to infection^25^. Bacteria counteract these effects by inducing SOS and DNA damage repair responses. In the present study, various genes encoding proteins involved in response to DNA damage (*priA, disA, dinG, mutT3, radA, recN, recC, recD, Rv0336, Rv0515* and *Rv3201c*) were upregulated. Additionally, genes involved in base excision repair, along with various DNA glycosylase enzymes to remove an altered base at the site of damage, endonucleases, polymerases and DNA ligase-encoding genes (*mutY, udgB, ung, Rv2191, ligD, dnaZX* and *Rv0142*) were also found to be upregulated.

Overall, these findings indicate that stressed mycobacteria within bone abscesses maintain their overall cellular integrity by synthesizing essential amino acids, strengthening their cell wall and repairing damaged DNA to withstand these detrimental effects.

### Downregulated pathways

The major pathways that were significantly enriched among the downregulated DEGs were protein metabolic processes, constituents of ribosomes, peptide biosynthesis and translational machinery (*p*≤0.001). Along with these, various other pathways, like cellular metabolic processes, organic substance biosynthesis (*p*≤0.005) and response to decreased oxygen level (*p*≤0.01) were negatively enriched, as mentioned in Table S5. The details of the major pathways and gene clusters are described below.

#### Translational machinery

Major downregulation of cellular protein metabolic processes, including structural constituents of ribosomes (*rpl, rpm* and *rps*), ribonucleoprotein complex, peptide biosynthetic processes (*valS, trpS, metS, lysS, ileS, glyS, cysS1* and *alaS*) and translational initiation and elongation factors (*rimP, frr, prfA, infB, infC, tsf, efp, tuf, typA, ideR* and *fiisA1*), was observed in the current study. Seventeen out of 22 *rps* genes (encoding the 30S ribosomal subunit) and 27 of 36 *rpl and rpm* genes encoding the 50S ribosomal subunit were downregulated (Table S4).

#### Protein excretion system

Along with major downregulation of the protein biosynthetic machinery, mycobacterial protein secretion systems were also downregulated in the current study. We found major downregulation of genes involved in type VII secretion systems, including *espl, espD, espA, espB, espC, espD, eccCa1, eccB1, esxA* and *esxB*. Additionally, 4 of the 8 *sec* genes (*secA1, secD, secE1* and *secF*) of the Sec secretion system and 1 of 4 *tat* genes (*tatA*), the twin-arginine (Tat) secretion system^9^ were downregulated (Table S5).

#### Mycobacterial growth

Several genes associated with *M. tuberculosis* cell division, including *ftsE, ftsH, ftsW, ftsX, ftsY and ftsZ*, encoding for septation-associated components of mycobacteria^26,27^ were significantly downregulated. Moreover, as shown in Table S5, 6 out of a total of 11 serine/threonine protein kinases (*pknA, pknB, pknD, pknE, pknI and pknL*), which are known to serve as environmental sensors regulating host-pathogen interactions and cell growth^28^ were found to be downregulated in this study. Additionally, we observed downregulation of 6 out of 7 *whiB* genes (*whiB1, whiB2, whiB3, whiB4, whiB6* and *whiB7*), transcriptional regulators suggestive of its role in mycobacterial growth and persistence^29^.

#### Respiratory machinery and ATP synthesis

Genes encoding several metabolic pathways associated with the growth of bacteria were found to be downregulated, including those involved in the *M. tuberculosis* respiratory pathway and energy generation. There was downregulation of genes coding for 6 subunits of NDH-I (*nuoA, nuoF, nuoK, nuoL, nuoM and nuoN*), cytochrome c oxidase (*ctaB, cta E, cta C and cta D*), cytochrome c reductase (*qcrA and qcrB*) which are involved in the routine respiratory pathway, as well as *ndhA* and *narGHJI*, which are involved in the alternative pathway of respiration. Also, there was downregulation of 4/8 subunits of genes encoding components of ATP synthase (*atpD, atpE, atpG* and *atpH*).

#### Cellular biosynthetic pathways

Cellular metabolism determines the fate of bacilli at the site of infection^30^. In this study, various biosynthetic pathways were repressed. Genes encoding cellular biosynthetic processes, like fatty acid biosynthesis (*kasA, kasB, accA, aacD1, accD5, accD6* and 20/34 total *fadD* genes), *de novo* synthesis of purines (*purA, purB, ndkA, nrdZ, nrdF1, nrdF2, guaA, guaB1, guaB3 and guaB2*), NAD biosynthesis (*nadA, nadB, nadE, nadD, gpm2, nudC, pncA and pncB2*), and biosynthesis of branched chain amino acids (*ilvA, ilvB1, ilvN, ilvB2, ilvC, ilvE, ilvD, leuD, leuA and leuB*) were downregulated. Along with these pathways, genes encoding proteins involved in maintaining the redox balance in mycobacteria were also downregulated. These included the genes encoding factors for the synthesis of: mycofactocin (*mftB, mftC, mftD, mftE and mftF*), a redox cofactor in mycobacteria; mycothiol biosynthesis (*mshA, mshB and mshD*), a glutathione analogue in mycobacteria; riboflavin (*ribA1, ribH, ribC and ribF*), a cofactor in redox system; and thioredoxin reductase (*trxB2*), which reduces thioredoxin, a redox protein. In addition, genes encoding NAD(P) transhydrogenases (*pntAa, pntAb and pntB*), which catalyzes the interconversion of NAD and NADP redox reaction, were also downregulated in the present study (Table S4).

#### Response to hypoxia

Mycobacteria may encounter different types of stress during host infection, including hypoxia within necrotic granulomas^31^. In order to survive such stresses, *M. tuberculosis* upregulates various genes. Interestingly, the vast majority of these hypoxia regulated genes (e.g., *devR, devS, groEL, groES, sigB, sigF and icl*) were found to be downregulated in bone TB lesions (Table S4).

Overall, our data reveal that there is downregulation of major metabolic processes essential for bacterial replication and growth, suggesting that *M. tuberculosis* enters a state of quiescence and reduced metabolism characterized by cell wall remodelling during chronic of infection within bone abscesses/necrotic tissue.

### qRT-PCR validation of microarray data

A small subset of genes (n=7) were randomly selected from microarray analysis which were further validated through qRT-PCR using the RNA isolated from the bone TB specimens. Out of the seven selected mycobacterial genes for validation, the gene expression of six DEGs was confirmed. Four of the genes (*Rv1230c, Rv2290 Rv1910c* and *Rv1971*) showed significant upregulation; *Rv3875* showed downregulation and *Rv1586* showed no change in the expression pattern similar to microarray (Fig. 1d). However, *Rv0986* was found to be upregulated by qRT-PCR with (log_2_ FC= 1.59±0.58) in contrast to the microarray analysis (log_2_Fc= −2.9), where it was downregulated.

### *In-silico* prediction of virulence factors

Seventeen of the mycobacterial genes showing >100-fold upregulation were further screened for identification of potential virulence factors with a significant role in bone TB pathogenesis.

Candidate virulence proteins were identified using Virulent Pred and VICM Pred. Five proteins were identified possessing virulence-like properties based on their amino acid composition, dipeptide composition, and PSI-BLAST and PSSM profiles. Two of the top upregulated genes, *Rv1046c* (conserved hypothetical protein) and *Rv1230c* (membrane protein), with log_2_fold changed of 9.7 and 9.1, respectively, along with *Rv3663c*, a dipeptide transporter (*DppD*), and two PE_PGRS family proteins, *Rv1441c* (*PE_PGRS26*) and *Rv2490c* (*PE_PGRS43*), were predicted to be virulence factors by *in silico* analysis (Table 2).

**Table 2:** Screening of mycobacterial virulence proteins encoded by genes amongst the top upregulated genes (FC>100) using Virulent Pred and VICM Pred. Proteins possessing virulence potential based on a minimum four features are highlighted in red color.

| S.No. | Rv ID | Virulent Pred |  |  |  |  |  |  | VICMPred |
| --- | --- | --- | --- | --- | --- | --- | --- | --- | --- |
|  |  | Amino acid Composition based | Dipeptide Composition Based | Similarity-Based using PSI-BLAST | PSI-BLAST created PSSM Profiles | Higher order Dipeptide Composition Based | Cascade of SVMs and PSI-BLAST | Virulent pred |  |
| 1 | **Rv1046c** | **Yes** | **Yes** | **No hits** | **Yes** | **Yes** | **Yes** | **Virulent** | **Metabolism Molecule** |
| 2 | **\*Rv1230c** | **Yes** | **Yes** | **Yes** | **No** | **No** | **Yes** | **Virulent** | **Cellular process** |
| 3 | Rv3086 | No | No | Yes | No | No | No | Non-Virulent | Metabolism Molecule |
| 4 | Rv2793c | No | No | No hits | No | No | No | Non-Virulent | Metabolism Molecule |
| 5 | Rv0775 | No | Yes | No hits | No | Yes | Yes | Non Virulent | Information and storage |
| 6 | Rv2844 | No | No | No hits | No | No | No | Non-Virulent | Cellular process |
| 7 | **Rv3663c** | **Yes** | **Yes** | **Yes** | **Yes** | **No** | **No** | **Virulent** | **Metabolism Molecule** |
| 8 | Rv2917 | No | No | No hits | No | No | No | Non-Virulent | Metabolism Molecule |
| 9 | **Rv2490c** | **Yes** | **Yes** | **Yes** | **Yes** | **Yes** | **Yes** | **Virulent** | **Information and storage** |
| 10 | **\*Rv1441c** | **Yes** | **Yes** | **Yes** | **Yes** | **Yes** | **Yes** | **Virulent** | **Information and storage** |
| 11 | Rv3721c | No | Yes | Yes | No | No | No | Non-Virulent | Information and storage |
| 12 | Rv2414c | No | No | No hits | No | No | No | Non-Virulent | Metabolism Molecule |
| 13 | Rv2736c | No | No | Yes | No | No | No | Non-Virulent | Cellular process |
| 14 | **\*Rv2672** | No | No | No hits | No | No | No | Non-Virulent | Cellular process |
| 15 | Rv3194c | No | No | Yes | No | No | No | Non-Virulent | Metabolism Molecule |
| 16 | Rv3667 | No | No | Yes | No | No | No | Non Virulent | Cellular process |
| 17 | Rv3500c | No | No | No | No | No | No | Non-Virulent | Cellular process |

### Confirmation of virulence gene expression in an in vitro model of bone TB

#### In vitro model of bone TB

In order to confirm the role of the in vivo expressed mycobacterial virulence genes in the pathogenesis of bone TB, an in vitro osteoblast cell line model was established by infecting the cells using *M. tuberculosis* H37Rv-lux (Fig. S1). *M. tuberculosis* established infection in osteoblasts, followed by an exponential increase in the bacterial burden up to 21days post-infection (Fig 3a). Osteoblast proliferation and alkaline phosphatase (ALP) activity was significantly decreased upon *M. tuberculosis* infection when compared to uninfected control cells at days 3 (p≤0.01), 7 (p≤0.05), 14 (p≤0.001) and 21 (p≤0.05) post-infection, as shown in Fig. S1.

**Fig. 3:**
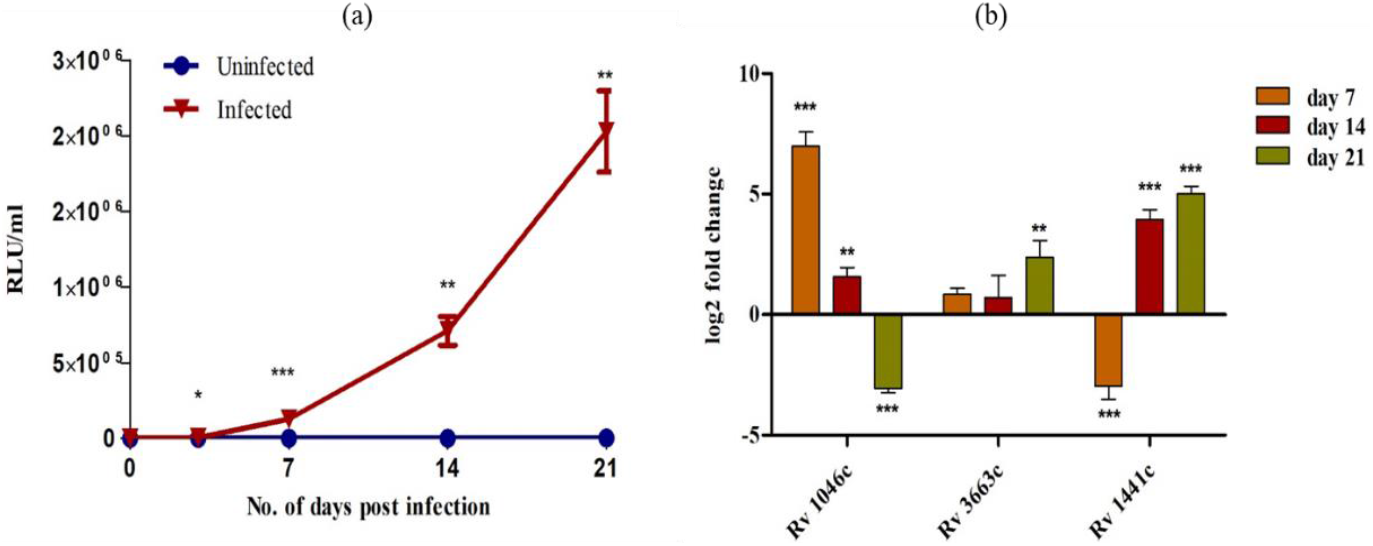
Intracellular multiplication of Mtb H37Rv-lux and gene expression analysis of intracellular Mycobacterium tuberculosis (i*M. tuberculosis*) within osteoblasts: MC3T3 osteoblast cell were infected with Mtb H37Rv-lux and incubated upto 21days post infection. a) Fold multiplication of Mtb in osteoblasts in terms of RLU/ml at different days of infection. p-value calculated by using student’s t-test to compare the infected vs. uninfected control samples at each time point. b) Gene expression analysis of in-silico identified virulent proteins of iMtb extracted from H37Rv-lux infected osteoblasts at different time points in comparison to H37Rv-lux. 16S rRNA was used as house-keeping gene for normalization. Log2 fold change was calculated using 2^^-ΔΔCt^. Each bar represents the mean ±SD of three different sets of experiment for each gene. One-way ANOVA with dunnett’s multiple comparison test was used to calculate statistical significance for gene expression of intracellular Mtb post 7d, 14d and 21d infection compared to control Mtb H37Rv-lux. *p<0.05, **p<0.01, ***p<0.001.

#### Gene expression analysis of mycobacterial virulence proteins in osteoblast cell culture

Gene expression profiling of virulence genes found to be upregulated in the in vivo bone TB lesions were further validated in the in vitro osteoblast model of bone TB using qRT-PCR. The gene *Rv1046c* showed significant upregulation at days 7 and 14 post-infection (p-value, ≤0.001), but was found to be downregulated by day 21 post-infection. *Rv1441c* showed significant downregulation at day 7 post-infection, although its expression increased afterwards and showed significant upregulation (p-value, ≤0.001) at days 14 and 21 post-infection. The gene *Rv3663* showed a significant time-dependent increase in gene expression post-infection as shown in Fig. 3b.

## Discussion

Bone is a highly mineralized tissue composed of bone-forming osteoblasts, bone degrading osteoclasts, osteocytes and extracellular matrix (ECM). Bone extracellular matrix (BEM) constitutes 30% of organic component majorly collagenous protein along with non-collagenous proteins and glycans; 70% of bone is made up of inorganic component (hydroxyapatite). BEM associated proteins play a key role in establishing infection through adherence, penetration and colonization by a pathogen within the bone. Many of the proteins such as collagen, bone sialoprotein, osteopontin and fibronectin are reported to be perfect niche for pathogens in the bone^32^. Bone TB primarily occurs through the dissemination of *M. tuberculosis* bacilli from the respiratory tract to the bones^33^. Enhanced bone resorption and bone loss are characteristic features of bone TB as a result of abnormal activation of osteoclasts^6,7,34^.

Although *M. tuberculosis* is known to cause bone TB, the cellular hosts for mycobacteria within the bone environment are still not well understood. Sarkar et al. demonstrated the replication of various mycobacterial strains in osteoblast cells^35^. Osteoblasts have also been shown to be host cells for *M. bovis* BCG infection^36^. As an intracellular pathogen, *M. tuberculosis* not only needs to survive in the host environment but also to replicate within human cells to disseminate. In order to establish infection in different tissues, *M. tuberculosis* is known to adapt by altering its transcriptional program in different environments^9,12^. To gain insight into the pathogenesis of bone TB, it is thus important to characterize the *M. tuberculosis* transcriptome in the bone microenvironment, particularly at the site of active disease. In the present study, we performed in vivo transcriptomic analysis of tubercle bacilli within abscesses or necrotic tissue obtained from patients with microbiologically-confirmed bone TB.

Overall, among DEGs, a greater number of genes were downregulated than were upregulated, corresponding to several metabolic pathways. Major significantly enriched upregulated pathways in human TB bone lesions were those involved in maintaining structural integrity and survival of the bacteria, including the synthesis of mAGP, LAM and glycolipids, which form the core structure of mycobacterial cell wall and are essential for *M. tuberculosis* resistance to various external stresses and reduced permeability to many drugs^37,38^. Likewise, two of the genes (*otsA* and *otsB2*) coding for trehalose synthesis and its transporter *mmpL13ab* were also significantly upregulated in this study. Induction of trehalose synthesis has been implicated in *M. tuberculosis* virulence, as it can be used as carbon and energy source during various stresses^39,40^.

In the current study, enrichment of several biosynthetic pathways of essential amino acids, including serine, arginine and cysteine, was observed. Serine biosynthesis is crucial for survival of mycobacteria inside the human host, as this amino acid is not taken up from the surrounding environment^41^. Besides acting as a nitrogen source, it is also important for the synthesis of other amino acids, such as glycine and cysteine. *De novo* synthesis of arginine is known to have a significant role in *M. tuberculosis* virulence, as arginine deprivation leads to accumulation of DNA damage, causing cell death^20,21^. The enzymes involved in this pathway are being considered as drug targets for TB therapeutics^42^. Thus, *M. tuberculosis* may upregulate genes involved in amino acid metabolism as a survival and/or pathogenesis strategy inside bone. Additionally, assimilation of inorganic nutrients, such as sulphur, in mycobacteria has been shown to contribute to mycobacterial virulence and survival^43^.

Mycobacterial protein phosphatases were also enriched in our study. *PstP* is an essential gene for *M. tuberculosis* intracellular survival and a key regulator of cell growth. Deletion of *pstP* causes cell wall defects leading to cell death, while overexpression of *pstP* lead to elongated cells with compromised cell survival^44^. We observed upregulation of *pknG*, which encodes an essential serine/threonine kinase responsible for sensing and responding to changes in nutrient availability^24^. Thus, the upregulated expression of these two regulatory genes, *pstP* and *pknG*, within bony TB lesions may be essential for metabolic switching from active growth to stasis in order to cope with environmental stresses. Along with these, mycobacterial genes encoding acid phosphatases enzymes (*Rv2577* and *Rv2135c*) were also found to be upregulated. Acid phosphatases secreted from osteoclast cells are considered a key marker of bone resorption, as they dissolve both organic (collagen) and inorganic (calcium and phosphorus) components of bone^45^. *Rv2577*, which encodes a secreted acid phosphatase^46,47^ containing a TAT (twin arginine translocation) motif was also found to be overexpressed, suggesting a potential role in bone resorption. Additionally, several genes (*mce1A, Rv3717, glmU* and *Rv0296c*) considered important for invasion of mycobacteria into host tissues were also induced in the present study.

One mechanism by which the host immune system responds to invading pathogens is the generation of various reactive oxygen species and reactive nitrogen intermediates, which cause damage to mycobacterial DNA^25^. Bacteria respond to such assaults by inducing DNA damage repair responses. In the present study, *M. tuberculosis* in bone lesions overexpressed genes involved in recombination repair systems (*recC, recD, recN* and *mutT3*) and base excision repair (*udgB, ung, Rv2191, dnaZX* and ligD). Induction of mycobacterial DNA damage repair responses promotes mycobacterial survival through adaptations to environmental stress and regulation of virulence^48^.

Amongst the negatively enriched pathways corresponding to downregulated genes in the study, major downregulation was observed in genes associated with cellular protein metabolism, growth of mycobacteria and several cellular biosynthetic processes. The bacteria surviving in bony lesions showed downregulation of ribosomal proteins involved in protein synthesis, which was accompanied by decreased protein export, as evidenced by downregulation of genes of mycobacterial secretion systems including *sec, tat* and several of the type VII secretion systems. The downregulation of *esxA* and *cfp10* from the Esx-1 secretion system is notable since the proteins encoded by these genes are considered important virulence factors of mycobacteria^49,50^. Other genes encoding Esat-6-like proteins were also downregulated in current study. The observed reduction in protein biosynthesis was accompanied by a downregulation of most genes encoding heat shock proteins, which are involved in proper protein folding during stress conditions.

Several metabolic pathways essential for mycobacterial growth were found to be downregulated in bone TB lesions. For example, the majority of purine biosynthesis genes, which are essential for *M. tuberculosis* growth^51,52^ were among the downregulated DEGs in the current study. As *M. tuberculosis* infection progresses from the acute to the chronic phase, immune activation and host-imposed stresses increase, which causes a metabolic shift in mycobacteria to ensure maintenance of membrane integrity in the absence of growth^30,53^. The genes involved in the synthesis of redox cofactors, such as NAD, mycofactocin, mycothiol, and riboflavin, which are important to resist oxidative stresses, were also found to be downregulated. Besides maintaining redox homeostasis, actively growing bacteria also needs energy in the form of ATP, which is synthesized by ATP synthase in response to the proton motive force generated through the respiratory chain. In the current study, the components of both the routine and alternative respiratory pathways, along with ATP synthase, were downregulated, thus indicating a low energy state of *M. tuberculosis* in the bone.

Furthermore, mycobacteria may come across different types of stresses during host infection, including hypoxia within necrotic granulomas and acidic pH in phagosomes^31^. In order to survive such stresses, *M. tuberculosis* upregulates various genes. A study by Rustad et al. has identified 49 genes, which are upregulated during *M. tuberculosis* exposure to hypoxia in vitro^13^. Interestingly, the great majority (40/49) of these hypoxia-regulated genes were found to be downregulated during bone TB. Another group of genes, which are upregulated during oxygen deprivation, include the universal stress response genes. Seven of 9 of these genes were downregulated in the current study. We also observed downregulation of 7/10 genes encoding heat shock proteins responsible for proper protein folding during stress conditions^54^. Downregulation of genes related to hypoxia and universal stress proteins points toward the diverse microenvironments encountered by mycobacteria within the lung parenchyma and bone. Thus, *M. tuberculosis* in bone lesions seems to be in a non-hypoxic, quiescent, low-energy state characterized by downregulation of genes involved in protein synthesis and transport, purine synthesis, and redox homeostasis.

After validating the microarray results using quantitative RT-PCR, the genes found to be the most highly upregulated in bone TB were further screened for the presence of *M. tuberculosis* virulence genes, with the hypothesis that these genes may play a significant role in the pathogenesis of bone TB. The bioinformatic tools VirulentPred and VICMPred, which have been used previously for such analyses^55,56^ identified several key virulence factors (Rv1046c, Rv1230c, dppD (Rv3663), PE_PGRS26 (Rv1440c) and PE_PGRS43 (Rv2490c)), which may play a role in *M. tuberculosis* pathogenesis and modulating host environment within the bone. Rv1046c is a hypothetical protein with unknown function belonging to a pathogenic genomic island (Rv1040c-Rv1046) with mobile genetic elements^57^, and *Rv1046c*-deficient mutants have been shown to have growth defects^52^. Rv1230c acts as cAMP-responsive stress regulator^58^. Rv3663, an oligopeptide/ dipetide permease (Opp)/Dpp transport system is involved in cell surface modulation of *M. tuberculosis*^59^. Expression of the Dpp transporters DppC and DppD in TB bone lesions could be important under nutrient-deficient conditions for the uptake of peptides^60^ from the extracellular matrix. Structural homology analysis showed that PE_PGRS 26 (Rv1441c) is an apoptosome-like protein associated with increased persistence of mycobacteria in mice and increased cell death and LDH release in macrophages^61^. Another PE-PGRS family of protein, Rv2490c has been shown to be expressed by *M. tuberculosis* in the lungs of guinea pig 30 and 90 days post-infection^62^.

Osteoblasts play a significant role and are key regulator in maintaining bone homeostasis. To study the importance of mycobacterial virulence proteins predicted in the present study in the pathogenesis of disease, an in vitro osteoblast cell line model was established with minor modifications from previous models^35,36^. Osteoblast cells were infected with *M. tuberculosis* H37Rv-lux in osteogenesis media for 21 days to allow the maturation of pre-osteoblasts to mature osteoblasts secreting extracellular matrix and minerals around osteoblasts^63^. The mycobacteria invaded the osteoblasts and multiplied exponentially, as determined in real time by relative luminescence intensity. Using RT-PCR in this model, we confirmed upregulation of bioinformatics-derived *M. tuberculosis* virulence genes, which were also found to be upregulated in human bone TB lesions, suggesting that the encoded proteins may play a role in the pathogenesis of bone TB.

### Concluding Remarks

The present study provides novel insights into mycobacterial adaptation in the bone microenvironment. Within human bone TB lesions during the chronic stage of infection, *M. tuberculosis* seems to be in a non-hypoxic, non-replicative and hypo-metabolic state characterized by alterations in its physiology, as reflected by decreased protein metabolism, export and cellular biosynthetic processes. In parallel, there is a major remodelling of cell wall synthesis favoring the maintenance of cellular integrity for bacillary survival within abscesses. Hence, it appears that after establishing a chronic infection within the bone, *M. tuberculosis* undergoes a transition from active growth to a metabolically quiescent state. These unique metabolic adaptations of mycobacteria during the chronic stage of infection can be further explored to gain an insight into mycobacterial pathogenesis and may lead to the development of novel therapeutic targets for the treatment of bone TB.

## Materials and Methods

### Study subjects and sample collection

To study the transcriptome of mycobacteria at the site of infection, abscess/necrotic tissue samples were used. Specimen were taken by an orthopaedic surgeon and collected in a sterile container from microbiologically confirmed (4 cases positive by GeneXpert MTB/RIF and MGIT culture and 1 case positive only by MGIT culture) bone TB patients before the commencement of anti-tubercular treatment (ATT).

Samples were collected only after obtaining the informed written consent from the patients visiting the Department of Orthopaedics, Post Graduate Institute of Medical Education and Research (PGIMER) in Chandigarh, India. The study was approved by the Institutional Ethics Committee vide no. PGI/IEC/2012/1334-35 and INT/IEC/2018/000126. The necrotic tissue/abscess samples were collected and immediately put into ice and then transferred to RNA later (Sigma Aldrich) and stored at −80°C till further use.

### Bacterial culture

*M. tuberculosis* H37Rv, a laboratory strain originally obtained from NCTC London, was grown in vitro in Sauton’s media supplemented with 10% OADC. A bioluminescent *M. tuberculosis* H37Rv strain (*M. tuberculosis-lux*), which stably expresses an integrated bioluminescent reporter (firefly *luxABCDE* full operon), including the luciferase enzyme and associated luciferin substrate (Dutta et al., 2020), was used for infection of osteoblast cells. *M. tuberculosis*-lux was grown in Middlebrook 7H9 broth containing 0.05%Tween, 0.2% glycerol and 10% OADC in a shaking incubator at 37°C and 200rpm.

### Cell line

The MC3T3 osteoblast cell line (ATCC CRL-2593) was used to establish an in vitro cell line model of bone TB. Osteoblast cells were maintained in α-MEM (Gibco) media containing 10% FBS (FBS; Corning) at 37°C in the presence of 5% CO_2_.

### RNA isolation

For isolation of *M. tuberculosis* RNA, stored samples were thawed on ice and centrifuged at 4500g for 15minutes to remove RNA later followed by two times of washing using chilled phosphate buffered saline (PBS). Further, the pellet obtained was treated with GTC solution (4M guanidium thiocyanate, 0.5% sarkosyl, 25mM tri sodium citrate, 0.1M β-mercaptoethanol and 0.5% Tween-80) for 5-10 minutes for effective removal of eukaryotic RNA, followed by centrifugation at 5000g for 20 minutes at 4°C and RNA was isolated using TRIzol reagent (Life Technologies), as described previously by Abhishek et al.^10^. RNA isolated from mid-logarithmic phase H37Rv culture was used as a reference to estimate the differentially expressed genes in bone TB specimens. Quality and quantity of RNA was determined using 2100 Bioanalyzer (Agilent) and Infinite 200 Pro NanoQuant (Tecan) respectively.

### Microarray

For microarray analysis, amplification and cyanine 3-CTP (Cy3) labelling of *M. tuberculosis* RNA was accomplished using One-Color Microarray-Based Low Input Quick Amp WT Labelling kit (Agilent Technologies) as per the manufacturer’s protocol using 300ng of input RNA. Further, labelled and amplified cRNA samples were purified using RNeasy Mini kit and quantified using Infinite 200 Pro Nano Quant plate spectrophotometer. cRNA concentration (ng/μL), 260/280 ratio and Cy3 concentration (pmol/μL) were measured to estimate the yield and specific activity of each sample. The samples (n=5) with a specific activity of more than 15 (pmol Cy3/μg cRNA) and yield of 0.825μg were selected and further processed for hybridization to customised *M. tuberculosis* array slides (custom GE array 8*15K; Agilent technologies G2509F-026323). 1.0 μg of labelled cRNA from each sample and control were used for co-hybridization using the gene expression hybridization kit (Agilent) as per the manufacturer’s protocol. The hybridized slides were washed and scanned using Sure Scan Microarray scanner (Agilent).

### Data extraction and analysis

Data were extracted from the scanned tiff image for each sample using feature extraction software and analysed using Gene Spring GX software (Agilent). For statistical analysis, student’s t-test was used with Benjamini-Hochberg’s correction. The genes with expression of >2 or ≤ −2 fold change and *p*-value ≤0.05 were filtered and considered to be significantly differentially expressed genes (DEGs). DEGs were subjected to functional categorization as per their functional categories listed in Tuberculist. Further, hypergeometric probability was used to find significantly enriched functional categories (*p*-value ≤0.05). Pathway enrichment analysis was done through Biocyc database (<u>BioCyc.org</u>) using fisher’s exact with Benjamini-Hochberg’s post hoc test.

### Validation of microarray results using real time qRT-PCR

Validation of microarray data was done using qRT-PCR. The RNA isolated from the clinical specimen from patients with confirmed bone TB was subjected to DnaseI (Thermo) treatment, followed by cDNA synthesis (BioRad iScript). The qRT-PCR was performed using Sybr Green master mix (Biorad) on the Rotor gene Q instrument (Qiagen). Relative gene expression of all the genes were calculated with the2^^(-ΔΔ*CT*)^ method, using 16S rRNA as an internal control and in vitro grown *M. tuberculosis* as reference. Primer sets used for relative gene expression are listed in the supplementary table (Table S1).

### Prediction of *M. tuberculosis* virulence proteins

Highly expressed genes with >100-fold upregulation were screened for prediction of genes encoding potential virulence proteins in bone TB. VICM Pred and Virulent Pred support vector machine (SVM)-based tools were used for prediction^55,56^.

### In vitro model of bone TB

Osteoblast cells were cultured in 24-well plates at a density of 5×10^4^ cells/well in osteogenesis media supplemented with 50μg/ml ascorbic acid and 2mM β-glycerophosphate for 24 hours at 37°C in the presence of 5% CO_2_. After 24 hours of incubation, osteoblast cells were infected with *M. tuberculosis*-lux at a MOI of 10-15 for 2 hours. Prior to infection, single-cell suspension of log-phase grown *M. tuberculosis-lux* (OD_600_ 0.3-0.5) was prepared followed by centrifugation, washing and vortexing in the presence of 3-mm glass beads. After 2 hours of infection, the monolayer of infected and uninfected control cells was washed thrice using PBS. Media containing amikacin (20ug/ml) was added to kill the extracellular bacteria, and then the cells were grown in amikacin-free osteogenesis media for 21 days post-infection. Culture media were changed every 2-3 days. Osteogenesis medium allows the differentiation of osteoblasts into mature osteoblast cells in the presence of mycobacteria. Cell proliferation was measured using the MTS assay (Promega), ALP activity using 1-step pNPP substrate solution (Thermo Scientific) and intracellular multiplication of bacteria was measured at days 0, 3, 7, 14 and 21 post-infection.

### Assessment of invasion and intracellular multiplication of mycobacteria

Mycobacterial burden within osteoblasts was measured in terms of relative luminescence units (RLU)/ml. At each time point, cells from were lysed using 0.1% TritonX-100 and collected in a micro-centrifuge tube. Immediately after collection, the cell lysate was pelleted and read for RLU using a luminometer (Promega GloMax 20/20).

### Gene expression analysis

qRT-PCR was used to study the relative gene expression of selected mycobacterial virulent proteins identified through microarray of bone TB patients in the intracellular *M. tuberculosis* isolated from osteoblast cell lines.

### Statistical analysis

For statistical analysis, Graph Pad Prism was used and the statistical difference between two groups was computed using unpaired Student’s t-test. For analysis of more than 2 groups, one-way ANOVA was used. Data were represented as mean ± standard deviation (SD). A p-value ≤ 0.05 was considered statistically significant.

## Acknowledgments

This work was supported by Indian Council of Medical Research (ICMR), project No.5/4-5/6/Ortho/2012-NCD-1. Training to KK arranged by Dr. Suman Laal under NIH/FIC training grant (1D43TW009588) is acknowledged. We also thank Mr. Yogesh Mittal for assistance in generating tables.

## References

1. Donoghue HD, Lee OYC, Minnikin DE, Besra GS, Taylor JH, Spigelman M. Tuberculosis in Dr Granville’s mummy: a molecular re-examination of the earliest known Egyptian mummy to be scientifically examined and given a medical diagnosis. Proc R Soc B. 2010;277(1678):51–6.

2. Kanade S, Nataraj G, Mehta P, Shah D. Pattern of missing probes in rifampicin resistant TB by Xpert MTB/RIF assay at a tertiary care centre in Mumbai. Indian Journal of Tuberculosis. 2019;66(1):139–43.

3. Rajasekaran S, Soundararajan DCR, Shetty AP, Kanna RM. Spinal Tuberculosis: Current Concepts. Global Spine Journal. 2018;8(4_suppl):96S–108S.

4. Jain AK. Tuberculosis of the spine: A fresh look at an old disease. The Journal of Bone and Joint Surgery British volume. 2010;92-B(7):905–13.

5. Jain A, Rajasekaran S. Tuberculosis of the spine. Indian J Orthop. 2012;46(2):127.

6. Tsumura M, Miki M, Mizoguchi Y, Hirata O, Nishimura S, Tamaura M, et al. Enhanced osteoclastogenesis in patients with MSMD due to impaired response to IFN-γ. Journal of Allergy and Clinical Immunology. 2021; S009167492100823X.

7. Hoshino A, Hanada S, Yamada H, Mii S, Takahashi M, Mitarai S et al. Mycobacterium tuberculosis escapes from the phagosomes of infected human osteoclasts reprograms osteoclast development via dysregulation of cytokines and chemokines. Pathogens and Disease. 2014; 70:28–39.

8. Jabir RA, Rukmana A, Saleh I and Kurniawati T. The Existence of Mycobacterium tuberculosis in Microenvironment of Bone. Mycobacterium - Research and Development, Wellman Ribón, IntechOpen. 2017. DOI: 10.5772/intechopen.69394.

9. Sharma S, Ryndak MB, Aggarwal AN, Yadav R, Sethi S, Masih S, et al. Transcriptome analysis of mycobacteria in sputum samples of pulmonary tuberculosis patients. PLOS ONE. 2017;12(3): e0173508.

10. Abhishek S, Saikia UN, Gupta A, Bansal R, Gupta V, Singh N, Laal S, Verma I. Transcriptional Profile of Mycobacterium tuberculosis in an in vitro Model of Intraocular Tuberculosis. Front Cell Infect Microbiol. 2018; 8:330.

11. Hudock TA, Foreman TW, Bandyopadhyay N, Gautam US, Veatch AV, LoBato DN, et al. Hypoxia Sensing and Persistence Genes Are Expressed during the Intragranulomatous Survival of Mycobacterium tuberculosis. Am J Respir Cell Mol Biol. 2017;56(5):637–47.

12. Rachman H, Strong M, Ulrichs T, Grode L, Schuchhardt J, Mollenkopf H, et al. Unique Transcriptome Signature of Mycobacterium tuberculosis in Pulmonary Tuberculosis. Infect Immun. 2006;74(2):1233–42.

13. Rustad TR, Harrell MI, Liao R, Sherman DR. The Enduring Hypoxic Response of Mycobacterium tuberculosis. PLoS ONE. 2018;3(1):e1502.

14. Betts JC, Lukey PT, Robb LC, McAdam RA, Duncan K. Evaluation of a nutrient starvation model of Mycobacterium tuberculosis persistence by gene and protein expression profiling: Nutrient starvation of M. tuberculosis. Molecular Microbiology. 2002;43(3):717–31.

15. Karp PD, Billington R, Caspi R, Fulcher CA, Latendresse M, Kothari A, et al. The BioCyc collection of microbial genomes and metabolic pathways. Briefings in Bioinformatics. 2019;20(4):1085–93.

16. Maitra A, Munshi T, Healy J, Martin LT, Vollmer W, Keep NH, et al. Cell wall peptidoglycan in Mycobacterium tuberculosis: An Achilles’ heel for the TB-causing pathogen. FEMS Microbiology Reviews. 2019;43(5):548–75.

17. Soni V, Upadhayay S, Suryadevara P, Samla G, Singh A, Yogeeswari P, et al. Depletion of M. tuberculosis GlmU from Infected Murine Lungs Effects the Clearance of the Pathogen. PLOS Pathogens. 2015;11(10):e1005235.

18. Haufroid M, Wouters J. Targeting the Serine Pathway: A Promising Approach against Tuberculosis. Pharmaceuticals (Basel). 2019;12(2):66.

19. Reitzer L. Amino Acid Synthesis: Reference Module in Biomedical Sciences. Elsevier; 2014. p. B9780128012383025000.

20. Tiwari S, van Tonder AJ, Vilchèze C, Mendes V, Thomas SE, Malek A, et al. Arginine-deprivation–induced oxidative damage sterilizes Mycobacterium tuberculosis. Proc Natl Acad Sci USA. 2018;115(39):9779–84.

21. Mizrahi V, Warner DF. Death of Mycobacterium tuberculosis by l-arginine starvation. Proc Natl Acad Sci USA. 2018;115(39):9658–60.

22. Khan MZ, Kaur P, Nandicoori VK. Targeting the messengers: Serine/threonine protein kinases as potential targets for antimycobacterial drug development. IUBMB Life. 2018;70(9):889–904.

23. Iswahyudi, Mukamolova GV, Straatman-Iwanowska AA, Allcock N, Ajuh P, Turapov O, et al. Mycobacterial phosphatase PstP regulates global serine threonine phosphorylation and cell division. Sci Rep. 2019;9(1):8337.

24. Rieck B, Degiacomi G, Zimmermann M, Cascioferro A, Boldrin F, Lazar-Adler NR, et al. PknG senses amino acid availability to control metabolism and virulence of Mycobacterium tuberculosis. PLoS Pathog. 2017;13(5): e1006399.

25. Vultos DT, Mestre O, Tonjum T, Gicquel B. DNA repair in Mycobacterium tuberculosis revisited. FEMS Microbiol Rev. 2009;33(3):471–87.

26. Datta P, Dasgupta A, Singh AK, Mukherjee P, Kundu M and Basu J. Interaction between FtsW and penicillin-binding protein 3 (PBP3) directs PBP3 to mid-cell, controls cell septation and mediates the formation of a trimeric complex involving FtsZ, FtsW and PBP3 in mycobacteria. Molecular microbiology. 2006;62: 1655–73.

27. Hett EC, Rubin EJ. Bacterial Growth and Cell Division: A Mycobacterial Perspective. Microbiol Mol Biol Rev. 2008;72(1):126–56.

28. Prisic S, Husson RN. Mycobacterium tuberculosis Serine/Threonine Protein Kinases. Microbiol Spectr. 2014;2(5):10.1128/microbiolspec.MGM2-0006-2013.

29. Zheng F, Long Q, Xie J. The Function and Regulatory Network of WhiB and WhiB-Like Protein from Comparative Genomics and Systems Biology Perspectives. Cell Biochem Biophys. 2012;63(2):103–8.

30. Warner DF. Mycobacterium tuberculosis Metabolism. Cold Spring Harbor Perspectives in Medicine. 2015;5(4): a021121–a021121.

31. Dutta NK, Karakousis PC. Latent tuberculosis infection: myths, models, and molecular mechanisms. Microbiol Mol Biol Rev. 2014;78(3):343–371. doi:10.1128/MMBR.00010-14

32. Hudson MC, Ramp WK, Frankenburg KP. Staphylococcus aureus adhesion to bone matrix and bone-associated biomaterials. FEMS Microbiology Letters. 1999;173(2):279–84.

33. Tuli S. Tuberculosis of the skeletal system. Fifth edition. New Delhi: Jaypee Brothers Medical Publishers; 2016.

34. Liu W, Zhou J, Niu F, Pu F, Wang Z, Huang M, et al. Mycobacterium tuberculosis infection increases the number of osteoclasts and inhibits osteoclast apoptosis by regulating TNF-α-mediated osteoclast autophagy. Exp Ther Med. 2020;20(3):1889–98.

35. Sarkar S, Dlamini MG, Bhattacharya D, Ashiru OT, Sturm AW, Moodley P. Strains of Mycobacterium tuberculosis differ in affinity for human osteoblasts and alveolar cells in vitro. SpringerPlus. 2016;5(1):163.

36. Hotokezaka H, Kitamura A, Matsumoto S, Hanazawa S, Amano S, Yamada T. Internalization of Mycobacterium bovis Bacillus Calmette-Guérin into osteoblast-like MC3T3-E1 cells and bone resorptive responses of the cells against the infection. Scand J Immunol. 1998;47(5):453–8.

37. Angala SK, Belardinelli JM, Huc-Claustre E, Wheat WH, Jackson M. The cell envelope glycoconjugates of Mycobacterium tuberculosis. Crit Rev Biochem Mol Biol. 2014;49(5):361–99.

38. Vincent AT, Nyongesa S, Morneau I, Reed MB, Tocheva EI, Veyrier FJ. The Mycobacterial Cell Envelope: A relict from the past or the result of recent evolution? Front Microbiol. 2018; 9:2341.

39. Thanna S, Sucheck SJ. Targeting the trehalose utilization pathways of Mycobacterium tuberculosis. Med Chem Commun. 2016;7(1):69–85.

40. Korte J, Alber M, Trujillo CM, Syson K, Koliwer-Brandl H, Deenen R, et al. Trehalose-6-Phosphate-Mediated Toxicity Determines Essentiality of OtsB2 in Mycobacterium tuberculosis In Vitro and in Mice. PLOS Pathogens. 2016;12(12):e1006043.

41. Borah K, Beyß M, Theorell A, Wu H, Basu P, Mendum TA, et al. Intracellular Mycobacterium tuberculosis Exploits Multiple Host Nitrogen Sources during Growth in Human Macrophages. Cell Reports. 2019;29(11):3580–3591.e4.

42. Mishra A, Mamidi AS, Rajmani RS, Ray A, Roy R, Surolia A. An allosteric inhibitor of Mycobacterium tuberculosis ArgJ: Implications to a novel combinatorial therapy. EMBO Mol Med. 2018;10(4).

43. Paritala H, Carroll KS. New Targets and Inhibitors of Mycobacterial Sulfur Metabolism. Infect Disord Drug Targets. 2013;13(2):85–115.

44. Sharma AK, Arora D, Singh LK, Gangwal A, Sajid A, Molle V, et al. Serine/Threonine Protein Phosphatase PstP of *Mycobacterium tuberculosis* Is Necessary for Accurate Cell Division and Survival of Pathogen. Journal of Biological Chemistry. 2016;291(46):24215–30.

45. Hannon RA, Clowes JA, Eagleton AC, Al Hadari A, Eastell R, Blumsohn A. Clinical performance of immunoreactive tartrate-resistant acid phosphatase isoform 5b as a marker of bone resorption. Bone 2004; 34:187–94.

46. Coker OO, Warit S, Rukseree K, Summpunn P, Prammananan T, Palittapongarnpim P. Functional characterization of two members of histidine phosphatase superfamily in Mycobacterium tuberculosis. BMC Microbiol. 2013; 13:292.

47. Forrellad MA, Blanco FC, Marrero Diaz de Villegas R, Vázquez CL, Yaneff A, García EA, et al. Rv2577 of Mycobacterium tuberculosis Is a Virulence Factor With Dual Phosphatase and Phosphodiesterase Functions. Front Microbiol. 2020;11:570794.

48. Bertok DZ. DNA Damage Repair and Bacterial Pathogens. Miller V, editor. PLoS Pathog. 2013;9(11):e1003711.

49. Guo S, Xue R, Li Y, Wang SM, Ren L et al. The CFP10/ESAT6 complex of Mycobacterium tuberculosis may function as a regulator of macrophage cell death at different stages of tuberculosis infection. Medical hypotheses. 2018; 78 (3):389–92.

50. Forrellad MA, Klepp LI, Gioffré A, Sabio y García J, Morbidoni HR, de la Paz Santangelo M, et al. Virulence factors of the Mycobacterium tuberculosis complex. Virulence. 2013;4(1):3–66.

51. Nours JL, Bulloch EM, Zhang Z, et al. Structural analyses of a purine biosynthetic enzyme from Mycobacterium tuberculosis reveal a novel bound nucleotide. J Biol Chem. 2011;286(47):40706–40716.

52. Sassetti CM, Rubin EJ. Genetic requirements for mycobacterial survival during infection. Proceedings of the National Academy of Sciences. 2003;100(22):12989–94.

53. Eoh H, Rhee KY. Multifunctional essentiality of succinate metabolism in adaptation to hypoxia in Mycobacterium tuberculosis. Proceedings of the National Academy of Sciences. 2013;110(16):6554–9.

54. Ryndak MB, Singh KK, Peng Z, Laal S. Transcriptional profiling of Mycobacterium tuberculosis replicating in the human type II alveolar epithelial cell line, A549. Genomics Data. 2015;5:112–4.

55. Garg, A., Gupta, D. VirulentPred: a SVM based prediction method for virulent proteins in bacterial pathogens. BMC Bioinformatics. 2008; 9:62.

56. Saha S, Raghava GPS. VICMpred: An SVM-based Method for the Prediction of Functional Proteins of Gram-negative Bacteria Using Amino Acid Patterns and Composition. Genomics, Proteomics & Bioinformatics. 2006;4(1):42–7.

57. Becq J, Gutierrez MC, Rosas-Magallanes V, Rauzier J, Gicquel B, Neyrolles O, et al. Contribution of horizontally acquired genomic islands to the evolution of the tubercle bacilli. Mol Biol Evol. 2007;24(8):1861–71.

58. Bai G, McCue LA, McDonough KA. Characterization of Mycobacterium tuberculosis Rv3676 (CRPMt), a cyclic AMP receptor protein-like DNA binding protein. J Bacteriol. 2005;187(22):7795–804.

59. Flores-Valdez MA, Morris RP, Laval F, Daffé M, Schoolnik GK. Mycobacterium tuberculosis modulates its cell surface via an oligopeptide permease (Opp) transport system. The FASEB Journal. 2009;23(12):4091–104.

60. Mitra A, Ko YH, Cingolani G, Niederweis M. Heme and hemoglobin utilization by Mycobacterium tuberculosis. Nat Commun. 2019; 10:4260.

61. Singh PP, Parra M, Cadieux N, Brennan MJ. A comparative study of host response to three Mycobacterium tuberculosis PE_PGRS proteins. Microbiology. 2008;154(11):3469–79.

62. Kruh NA, Troudt J, Izzo A, Prenni J, Dobos KM. Portrait of a Pathogen: The *Mycobacterium tuberculosis* Proteome In Vivo. PLOS ONE. 2010;5(11): e13938.

63. Buttery LD, Bourne S, Xynos JD, Wood H, Hughes FJ, Hughes SP, et al. Differentiation of osteoblasts and in vitro bone formation from murine embryonic stem cells. Tissue Eng. 2001;7(1):89–99.

